# GPR17 is an Essential Component of the Negative Feedback Loop of the Sonic Hedgehog Signalling Pathway in Neural Tube Development

**DOI:** 10.1101/424598

**Authors:** Atsuki Yatsuzuka, Akiko Hori, Minori Kadoya, Mami Matsuo-Takasaki, Toru Kondo, Noriaki Sasai

**Affiliations:** Developmental Biomedical Science, Graduate School of Biological Sciences, Nara Institute of Science and Technology, 8916-5, Takayama-cho, Ikoma 630-0192, Japan; Department of Regenerative Medicine and Stem Cell Biology, Faculty of Medicine, University of Tsukuba, Tsukuba 305-8575, Japan; Division of Stem Cell Biology, Institute for Genetic Medicine, Hokkaido University, Kita-15, Nishi-7, Kita-Ku, Sapporo 060-0815, Japan; Present Address: iPS Cell Advanced Characterization and Development Team, RIKEN BioResource Research Center, 3-1-1 Koyadai, Tsukuba, Ibaraki 305-0074, Japan

## Abstract

Dorsal-ventral pattern formation of the neural tube is regulated by temporal and spatial activities of extracellular signalling molecules. Sonic hedgehog (Shh) assigns ventral neural subtypes via activation of the Gli transcription factors. Shh activity changes dynamically during neural differentiation, but the mechanisms responsible for regulating this dynamicity are not fully understood. Here we show that the P2Y-type G-protein coupled receptor GPR17 is involved in temporal regulation of the Shh signal. GPR17 was expressed in the ventral progenitor regions of the neural tube and acted as a negative regulator of the Shh signal in chick embryos. While the activation of the GPR17-related signal inhibited ventral identity, perturbation of *GPR17* expression led to aberrant expansion of ventral neural domains. Notably, perturbation of *GPR17* expression partially inhibited the negative feedback of Gli activity. Moreover, GPR17 increased cAMP activity, suggesting that it exerts its function by inhibiting the processing of Gli3 protein. GPR17 also negatively regulated Shh signalling in neural cells differentiated from mouse embryonic stem cells, suggesting that GPR17 function is conserved among different organisms. Our results demonstrate that GPR17 is a novel negative regulator of Shh signalling in a wide range of cellular contexts.

**Author Summary:** During neural development, determination of cell fate and the progress of differentiation are regulated by extracellular signal molecules, including Sonic Hedgehog (Shh). Shh forms a gradient within the embryonic organ of the central nervous system, or the neural tube, and a variety of cells are produced corresponding to the concentration. While the signal concentration is critical for cell fate, recent studies have revealed that the intracellular signal intensity does not always correspond to the Shh concentration. Rather, the intracellular signal intensity changes over time. Importantly, the signal intensity peaks and gradually decreases thereafter, and the half-life of the Shh signal contributes to the cell fate determination. However, the mechanisms for this temporal change are not fully understood.

By using chick embryos and mouse embryonic stem cells as model systems, we demonstrate that the G-protein coupled receptor, GPR17, is an essential regulator for the negative feedback of the Shh signal during neural development. While *GPR17* gene expression is induced by the Shh signal, GPR17 perturbs the Shh signalling pathway. This negative function of GPR17 on the Shh signal is conserved among different vertebrate species. The collective data demonstrate that GPR17 is a negative regulator for the Shh signalling pathway in a wide range of the cellular contexts.

## Introduction

The neural tube, the embryonic organ of the central nervous system, consists of neural progenitors and post-mitotic neurons arrayed in an orderly manner [1,2]. During neural tube development, extracellular signalling molecules (“morphogens”) produced in the signalling centres of the organ provide positional cues to uncommitted neural progenitor cells, and thereby assign the cellular fates in a concentration-dependent manner [3]. Identifying the molecular mechanism by which cells convert positional information into fate determination is one of the major goals of developmental biology.

Sonic hedgehog (Shh) is a signalling molecule expressed in the floor plate (FP) of the neural tube and its underlying mesodermal tissue, the notochord. Shh plays essential roles in the assignment of ventral identities [3]. At the cellular level, Shh ligand binds to the 12-span transmembrane protein Patched 1 (Ptch1) at the primary cilia. The binding activates another membrane protein, Smoothened (Smo). Smo further activates the transcription factors Gli2/3, which are transported into the nucleus to induce the expression of their target genes [3]. Gli2 and Gli3 have dual activities; in the absence of Shh, the precursor Gli2/3 proteins associate with the scaffold protein SuFu to form complexes with Cullin3 and Protein Kinase A (PKA), and are phosphorylated and proteolytically processed into their repressor forms [4-7]. In response to the Shh ligand, these complexes are dissociated, ubiquitination is perturbed, and Gli2/3 are converted into their active forms [7-9].

In the spinal cord, Shh forms a ventral-to-dorsal gradient with the highest level in the FP, and is required for the assignment of at least five ventral neural progenitor domains in addition to FP [1,3]. The ventral domains include p0-p2, pMN, p3 that are arrayed in this order from dorsal to ventral, and can be defined by transcription factors expressed in a domain-specific manner. For instance, two transcription factors Olig2 and Nkx2.2 are expressed in the pMN and p3 domains, respectively, and are essential for these identities [10-12]. During the neural tube development, Olig2 responds to Shh and is expressed first, and the expression of Nkx2.2, which is also a responsive gene to Shh, expression begins later [13]. The higher concentration of Shh allows the quicker switch of the progenitor cells from Olig2- to Nkx2.2-expressing state [14]. In this manner, the progenitor cells exposed to a higher Shh concentration form the Nkx2.2-positive p3 domain at a more ventral position than the Olig2/pMN domain. Thus, the concentration of Shh is converted into the dynamic expression of the transcription factors and determines the cell fates in the spinal cord [15].

The Gli activity partially reflects the concentration of Shh at the initial step of neural differentiation [14]. After it is initially elevated, however, Gli activity decreases over time [14,15]. Moreover, the half-life of the signal intensity determines the kinetics of the conversion of the downstream transcription factors that characterise the neural domains. Therefore, the dynamic change in Gli activity, which is termed temporal adaptation, is important for correct pattern formation of the neural tube [13-16].

Multiple mechanisms have been proposed to explain this adaptation [17]. One model involves upregulation of the *Ptch1* gene. Since Ptch1 is a negative regulator of the Shh signal, accumulation of Ptch1 depletes Shh and consequently decreases its intracellular signalling activity [13]. Another model proposes that the temporal decrease in the level of Gli2, which is a mediator protein of the Shh signal, decreases over time, suggesting that the transduction of the Shh signal is reduced as the neural development proceeds [17]. Importantly, this adaptation takes place only in the context of the neural differentiation, and not in cultured cells, such as NIH3T3 fibroblast cells [1,17,18], suggesting that some key genes are specifically induced in a developmental context. However, such regulators have yet to be identified.

In parallel with Shh activity, intracellular cyclic AMP (cAMP) level and the activity of the downstream mediator PKA have critical roles for the neural tube pattern formation [5,19]. Absence of PKA activity affects the subcellular localisation and processing of Gli proteins at the cellular level, and results in the expansion of the ventral domains of the neural tube [5]. Likewise, the activity of adenylyl cyclase 5 and 6 (AC5 and AC6, respectively), which encourage the production of cAMP by resolving adenosine triphosphate (ATP), affects the Gli activity and determination of the ventral cell fate [20,21]. Moreover, G-protein coupled receptors (GPCRs) have been suggested to control the neural tube pattern formation through regulating the intracellular cAMP level [18,22]. GPCRs couple with G-proteins that are comprised of Gα, Gβ and Gγ subunits. G-proteins are categorised into several subclasses based on the type of the Gα protein; Gα_s_ and Gα_q_ can potentiate the cAMP level, whereas Gα_i_ decreases this level [23]. With respect to the neural tube development and Shh signal, GPR161 has been suggested to interact with Gα_s_ and negatively regulates the ventral neural identities by elevating the cAMP level [24]. Conversely, GPR175 interacts with Gα_i_ protein and enhances the Shh signal [25]. Therefore, the GPCR/cAMP/PKA axis is a critical regulatory pathway in the neural tube pattern formation. However, the mechanisms by which cAMP/PKA activity is involved in the temporal changes of Gli activity remain to be revealed.

In this study, we hypothesised the existence of a GPCR regulated by the Shh signal. To identify GPCRs induced by Shh, we performed a quantitative RT-PCR (RT-qPCR)-based screen in chick neural explants, and identified *GPR17* as a candidate gene. *GPR17* is induced by Shh, but negatively regulates the Shh signal, making it an example of negative feedback. GPR17 also functions as a negative regulator in neural differentiation of mouse embryonic stem (ES) cells. Taken together, our findings demonstrate that GPR17 is a negative regulator in multiple cellular contexts.

## Results

### GPR17 is expressed in the ventral region of the neural tube

To identify genes involved in Shh signal dynamics, we focused on GPCRs, as many GPCRs regulate the activity of PKA and further modify the processing of Gli transcription factors, their activities [21]. With this in mind, we selected genes of GPCRs that can bind to Gα_s_ or Gα_q_, and performed a screen that combined RT-qPCR and in situ hybridisation (S1 Fig) to isolate the GPCR that responds to Shh. Based on its expression pattern and responsiveness to Shh, we identified the P2Y-like GPCR (a group of GPCRs that take purine as ligand) *GPR17* as a candidate. *In situ* hybridisation analysis revealed that *GPR17* is expressed broadly in the ventral region of the trunk neural tube at Hamburger-Hamilton (HH) stage 14 (Fig 1A). At HH stage 24, the expression formed a dorsal-to-ventral gradient in the progenitor region of the neural tube, with the lowest level at the dorsal region and highest level at Olig2-positive pMN and Nkx2.2-positive p3 regions, which are motor neuron and V3 interneuron progenitor regions, respectively (Fig 1B and 1C) [2]. In addition, while the *GPR17* expression was found in a wide area in the ventral at the beginning of the neural tube development (Fig 1A), the floor plate expression was abolished as development progressed (Fig 1B and 1C). In addition, while the expression of *GPR17* occurred in a wide area in the ventral region at the beginning of the neural tube development (Fig 1A), the floor plate expression halted as development progressed (Fig 1B and 1C).

**Figure 1:**
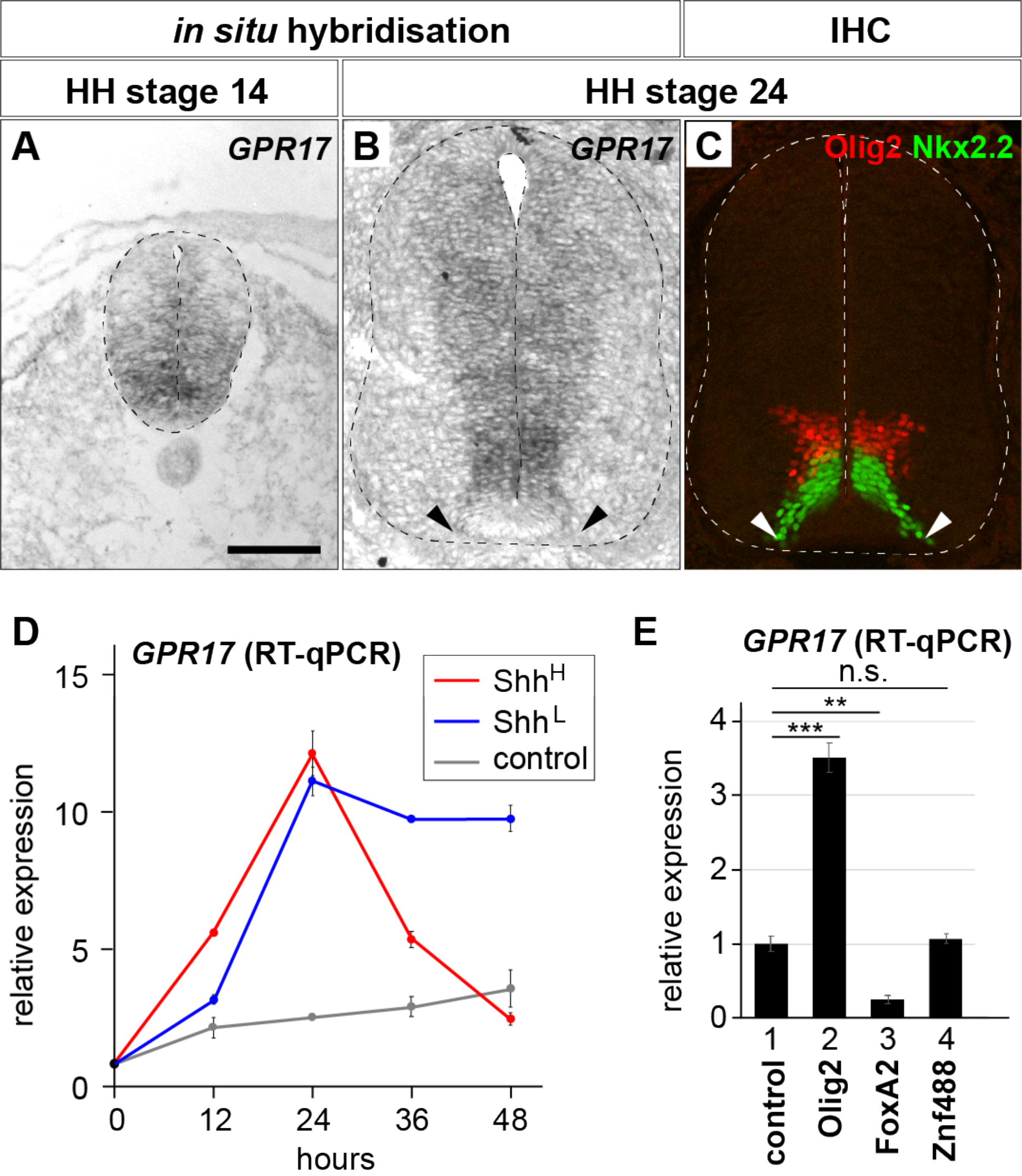
GPR17 is strongly expressed in ventral progenitor regions of the neural tube. (**A**-**C**) *In situ* hybridisation analysis of *GPR17* on a section of trunk neural tube was performed at HH stage 14 (**A**) and 24 (**B**), and the expression pattern was compared with those of Olig2 (red) and Nkx2.2 (green), as determined by immunohistochemistry (IHC) (**C**). The ventral ends of the *GPR17* (**B**) and Nkx2.2 (**C**) expression are indicated by arrowheads. In (**B**) and (**C**), expression analyses were performed on the adjacent sections. (**D**) *GPR17* was induced by the Shh signal, and expression was maintained when the explants were treated with Shh^L^, whereas those treated with Shh^H^ lost expression after being upregulated. Neural explants were treated in the absence or in the presence of Shh^L^ or Shh^H^ for the indicated time periods and analysed by RT-qPCR. (**E**) The *GPR17* expression is affected by Olig2 and FoxA2, but not by Znf488. Neural explants electroporated with *control GFP*, *Olig2*, *FoxA2* or *Znf488* were prepared and the expression of GPR17 was analysed by qPCR after the 48 hour-culture. Scale Bar in **A** = 100 μm, **B**,**C** = 50 μm. **: p<0.01, ***: p < 0.001, n.s.: not significant.

To investigate the relationship between Shh and *GPR17* expression, we performed an *ex vivo* analysis using intermediate neural explants. We isolated neural progenitor cells from the preneural tube area [26] of HH stage 9 embryos and cultured the cells for 48 hours in the presence of low or high concentrations of Shh (Shh^L^ and Shh^H^, respectively), which determine motor neuron and floor plate identities, respectively (S2 Fig). Expression levels of *GPR17* were measured by RT-qPCR, every 12 hours up to 48 hours (Fig 1D). At 12 hours, *GPR17* expression was higher in explants treated with Shh^L^ or Shh^H^ than in untreated explants (Fig 1D), and this trend continued at 24 and 36 hours. By contrast, at 48 hours, expression was downregulated in explants treated with Shh^H^, but remained high in explants treated with Shh^L^. This finding is consistent with the *in vivo* expression pattern of abolished expression in the floor plate (Fig 1A).

Next we explored the upstream transcription factors that induce *GPR17* expression. Since *GPR17* requires *Olig1* for its expression [27] and is one of the target genes of Olig2 during oligodendrocyte development [28], we assessed if the overexpression of an Olig-type transcription factors could induce the *GPR17* transcript. To address this, we prepared intermediate neural explants that were electroporated with the *Olig2* expression plasmid, and determined the *GPR17* expression level by RT-qPCR. *GPR17* expression was elevated at 24 hours, suggesting that Olig2 alone is sufficient to induce *GPR17* expression (Fig 1E, lanes 1 and 2). By contrast, FoxA2, which is expressed in the floor plate, downregulated *GPR17* (Fig 1E, lanes 1 and 3), which was consistent with the observation that the *GPR17* expression was lower in the floor plate region. Znf488 (also known as Zfp488), which is expressed in the same regions as *Olig2* [29,30], did not have a significant effect on GPR17 expression (Fig 1E, lanes 1 and 4). These data supported the idea that *GPR17* expression is targeted by Olig2 [28], and becomes exclusive to the floor plate as development progresses.

The collective results suggested that GPR17 expression is mainly regulated by Shh signal and its downstream transcription factor Olig2. This intriguing regulation of *GPR17* expression prompted us to further investigate its role in neural tube development.

### GPR17 is a negative regulator of the Shh signalling pathway

We sought to characterise the molecular role of GPR17 in the Shh signalling pathway. For this purpose, we first analysed the subcellular localisation of GPR17 by immunohistochemistry in NIH3T3 cells (Fig 2A-2D’). GPR17 was detected throughout the cells (Fig 2B) when GPR17 primary antibody was added (Fig 2A and 2B). The distribution of the GPR17 signal did not change upon treatment with Shh (Fig 2C) or its agonist SAG (Fig 2D) [31]. Importantly, while the intracellular signal transduction induced by Shh is mediated by the cilia, GPR17 was not significantly localised to the ciliary shaft or its surrounding areas in any conditions (Fig 2A’, 2B’, 2C’, 2D’).

**Figure 2:**
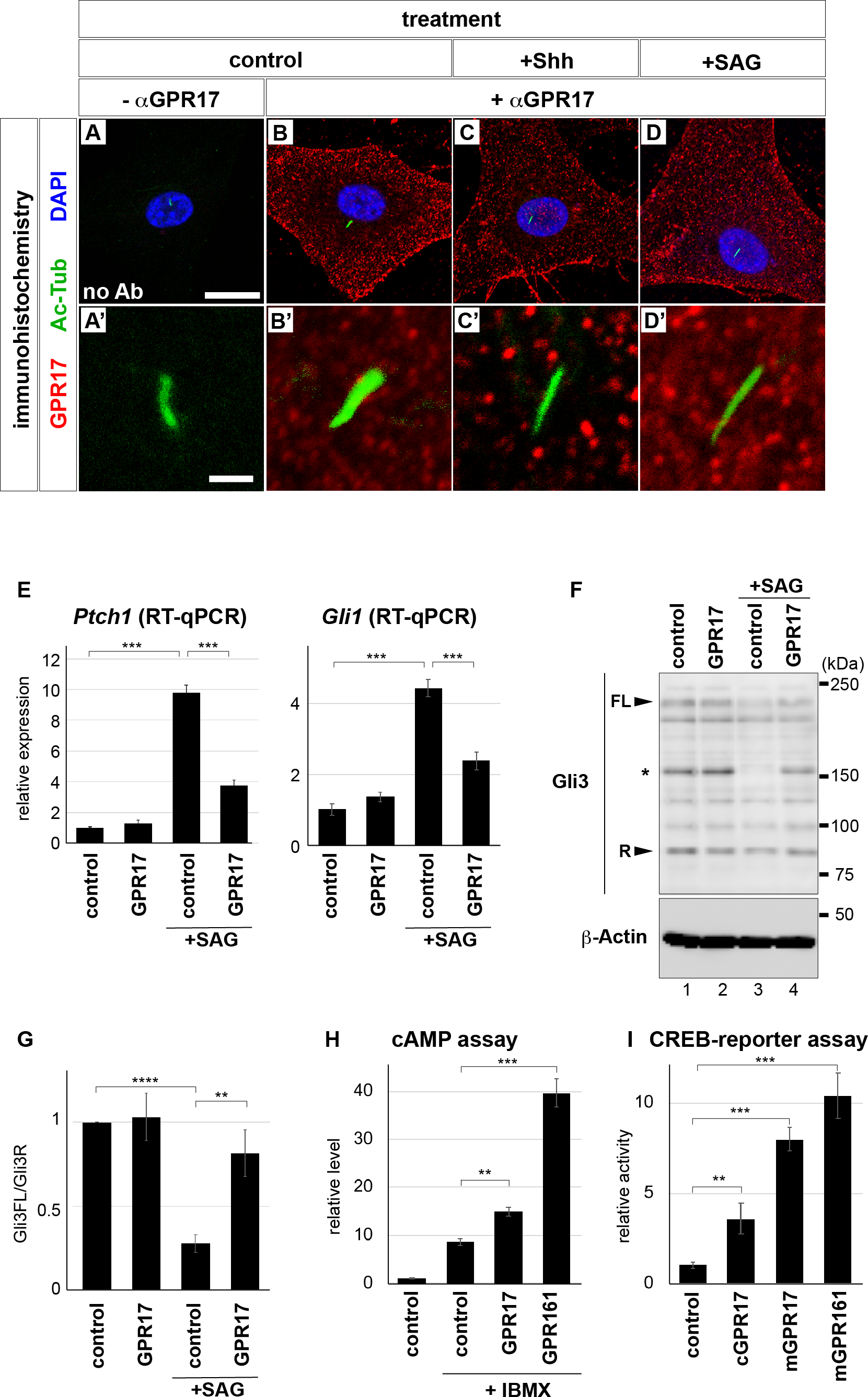
GPR17 is a negative regulator for the Shh signalling pathway. (**A**-**D’**) Subcellular localisation of GPR17. NIH3T3 cells were incubated without (**A**-**B’**) or with Shh^H^(**C,C’**) or 500 nM of SAG (**D,D’**). Immunohistochemistry was performed without (**A,A’**) or with the GPR17 antibody (**B-D’**) in addition to the Ac-Tub antibody. (**E**) Overexpression of GPR17 decreased the expression levels of *Ptch1* and *Gli1* following SAG treatment. NIH3T3 cells transfected with *YFP* or *GPR17-YFP* were treated with SAG for 24 hours, and expression of *Ptch1* and *Gli1* was analysed by RT-qPCR. (**F**) The ratio of Gli3FL and Gli3R was altered in the *GPR17-YFP*-transfected cells. Control *YFP* (lanes 1,3) or *GPR17-YFP* (lanes 2,4) expression plasmids were transfected into the NIH3T3 cells and cells were cultured with SAG for 6 hours. The cell lysates were analysed by western blotting. The bands indicated by an asterisk (*) always appear but are apparently not Gli3. (**G**) The quantitative data of Gli3FL/Gli3R for (**F**). (**H**) The cAMP level is upregulated by GPR17. NIH3T3 cells were transfected with control *YFP* or *GPR17-YFP* expression plasmids. 3-isobutyl-1-methylxanthine (IBMX) was added at 1 μM in the last 30 minutes. Intracellular cAMP was measured. (**I**) The CREB activity was measured by a luciferase assay. Scale Bar in **A** for **A,B,C,D** = 20 μm and in **A’** for **A’,B’,C’,D’** = 2 μm. **: p<0.01, ***: p < 0.001.

Next, to examine the effect of GPR17 in the Shh intracellular signalling pathway, cells transfected with either yellow fluorescent protein (YFP) or GPR17-YFP were treated with SAG for 24 hours. RT-qPCR analysis was conducted to assay the expression of *Ptch1* and *Gli1*, which are primary target genes of the Shh signal [17] (Fig 2E). The extent of the upregulation of these genes in *GPR17-*transfected cells was significantly lower than in the control cells (Fig 2E). Given that SAG targets the Smo protein [31], this result suggests that GPR17 perturbs the intracellular Shh signal downstream of Smo.

We therefore assessed if the reduction of the target genes of the Shh signal resulted at the step of Gli processing by examining the accumulation of Gli3 protein in cells. Gli3, a critical mediator of the Shh signalling pathway [32], is a dual-mode transcription factor with the ubiquitinated and truncated repressor form and the full-length form that can be active by further modifications [7,8,33,34]. Moreover, it has been suggested that the altered ratio of the amounts of the full-length and repressor form, as well as the total amount of the protein, changes upon the exposure to Shh or its related signals [18]. Therefore, we attempted to determine whether this ratio was affected by the overexpression of *GPR17*. We prepared control-*YFP* or *GPR17-YFP*-transfected cells that were untreated or treated with SAG and examined them by western blots using the Gli3 antibody (Fig 2F). Without SAG treatment, the ratio of the full-length over the repressor form of Gli3 was not changed by the transfection of GPR17 (Fig 2F, lanes 1 and 2). By contrast, when cells were treated with SAG, Gli3FL became less abundant while the amount of Gli3R was less affected, consistent with the previous report that Gli3 is destabilised by the Shh signal [18] (Fig 2F, lane 3 and Fig 2G). However, in the GPR17-YFP-expressing cells, the change in Gli3FL was less than that of the control cells (Fig 2F, lane 4 and Fig 2G), suggesting that the effect of SAG on Gli3 was perturbed by the presence of GPR17.

We further measured the cAMP levels in the cells, because modification of Gli3 proteins depends on the intracellular cAMP level [6]. Cells were transfected with control vector or the plasmid conveying *cGPR17* (chicken *GPR17*) and treated with 3-isobutyl-1-methylxanthine (IBMX), a non-competitive selective phosphodiesterase inhibitor, to raise the basal level of intracellular cAMP (Fig 2H). *GPR161* was used as a positive control [24]. A higher cAMP level was evident in GPR17 cells than in control cells (Fig 2H). To further support the idea that the cAMP level was upregulated in the GPR17-expressing cells, we conducted a reporter assay measuring the activity of the cAMP-responsive element-binding region (CREB). Transfection of both *cGPR17* and *mGPR17* (mouse GPR17) expression plasmids raised the CREB activity, confirming that the intracellular cAMP level was raised upon the overexpression of *GPR17* (Fig 2I).

Taken together, these findings demonstrated that GPR17 functions as a negative regulator of the Shh signalling pathway by upregulating cAMP levels.

### GPR17 negatively regulates the ventral identity of the neural tube

To investigate the activity of GPR17 in neural tube development and pattern formation, we overexpressed *GPR17* by *in ovo* electroporation in the neural tube of the chick embryos, and monitored its effect on dorsal-ventral pattern formation. However, pattern formation was not significantly altered (0/7 embryos; S4A-S4C” Fig). This could be because a trigger, such as a ligand, is necessary to activate GPR17-mediated signalling, with the overexpression of *GPR17* alone not being sufficient to fully activate the receptor.

Therefore, we speculated that the combination of GPR17 with its specific agonist, MDL29951 [35,36], would embody the phenotype of the *GPR17* overexpression, and attempted to administrate the chemical on the embryos. To administer MDL29951 to embryos, we employed New culture system, in which embryos were cultured from HHs stage 12 *ex ovo* to maintain the concentration of the chemical throughout the culture [37] (Fig 3A-3D). In the control condition, the embryos displayed expression of Olig2 (Fig 3A) and Pax7 (Fig 3C) in the ventral and dorsal regions, respectively, after 36 hour-culture. By contrast, when the embryos were cultured in the presence of MDL29951, the expression of Olig2 was reduced (Fig 3B) and Pax7 was expanded ventrally (Fig 3D), suggesting the dorsal-ventral pattern was affected by the compound.

**Figure 3:**
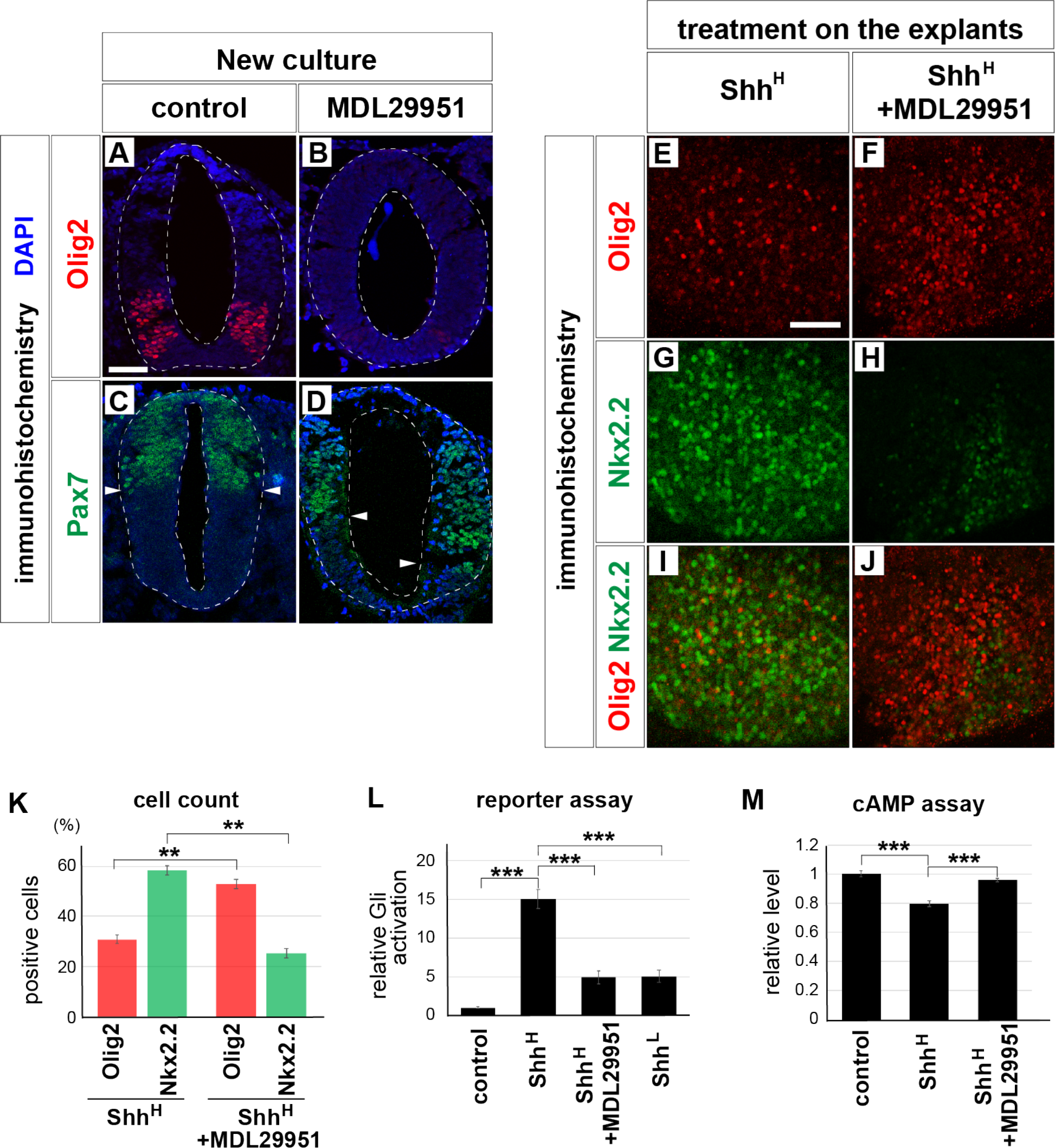
GPR17 negatively regulates the Shh signalling pathway and affects the cell identity of neural progenitors. (**A**-**D**) The ventral identity was affected by MDL29951. The HH stage 12 embryos were taken out from the eggs, and were cultured for 36 hours with the New culture system containing the control solvent (DMSO; **A,C**) or 100 μM of MDL29951 (**B,D**). (**E**-**J**) Ventral identities are altered by treatment with MDL29951, a specific agonist of GPR17. Explants were incubated for 24 hours with Shh^H^ in the absence (**E,G,I**) or presence (**F,H,J**) of 30 μM MDL29951, and stained with Olig2 (**E**,**F**,**I**,**J**) and Nkx2.2 (**G**,**H**,**I**,**J**) antibodies. Merged images in (**I**,**J**) and quantification in (**K**).(**L**) Gli activity is reduced in the neural explants treated with MDL29951 for 24 hours. Explants electroporated with GBS-Luc were treated with control medium, Shh^H^, Shh^H^ + MDL29951, or Shh^L^. (**M**) cAMP level is upregulated by the treatment with MDL29951 in the intermediate neural explants. Explants were prepared and treated with control, Shh^H^ or Shh^H^ + MDL29951 for 24 hours. IBMX was added at 1 μM for the last 1 hour. Scale Bars in **A** for **A**-**D**, **E** for **E**-**J** = 50 μm.

Next, in order to reveal the relationship between the intracellular Shh signal and GPR17 more directly, we investigated the effect of GPR17 and MDL29951 on the forced activation of the Shh signal. We electroporated the constitutively-active form of Smo (*SmoA1*) [38] at HH stage 12 and found that the neural tube was highly ventralised at 48 hours post transfection (hpt), with a strong expansion of the Nkx2.2-positive region and the repression of the Pax7-positive area (S3A-S3B”, S3FS3G” Fig). The administration of MDL29951 to the *SmoA1*-electroporated embryos (S3C-S3C”, S3HS3H” Fig) did not change the trend, suggesting that the concentration of MDL29951 used was not sufficient to influence the Shh signal activated by Smo. The co-electroporation of *GPR17* with *SmoA1* also did not change the expansion of the ventralisation (S3D-S3D”, S3I-S3I” Fig). However, when the same amount of MDL29951 was administered to *SmoA1* and *GPR17* electroporated neural tubes, the extent of the ventral expansion was significantly reduced (S3E-S3E” Fig), and the number of electroporated cells expressing Pax7 increased (S3J-S3J”, S3K Fig). The results suggest that MDL29951 synergistically works with GPR17 and that the activation of GPR17-mediated signal perturbs the intracellular Shh signalling pathway induced by Smo. Moreover, it is unlikely that the alteration of the pattern formation is mediated by the programmed cell death, as a TdT-mediated dUTP nickend labelling (TUNEL) assay did not detect a significantly increased number of positive signals (S3L-S3M’ Fig).

We further investigated the direct effect of GPR17 on the Shh signalling pathway by an intermediate neural explant analysis. Because the *GPR17* expression was induced by Shh at 24 hours in the explants (Fig 1D, S1 Fig), we expected that the treatment with MDL29951 on the explants would alter the neural identities. Shh^H^ treatment of the explants for 24 hours simulated the differentiation into ventral neural progenitor cells. Nkx2.2 expression was detectable in the majority of cells, and Olig2 in a smaller subset (4 areas; Fig 3E, 3G 3I, 3K). By contrast, when the explants were treated with MDL29951 along with Shh^H^, the number of cells expressing Nkx2.2 decreased, whereas the number of Olig2-positive cells increased (4 areas; Fig 3F, 3H, 3J, 3K). Given that the Olig2-expressing cells can be induced by Shh^L^, whereas Nkx2.2 expression is induced by Shh^H^ in 24-hour cultures [13], the results suggest that Shh was partially inhibited by the GPR17-associated signalling. This effect occurred independently from the programmed cell death, as the increasing number of TUNEL-positive cells was not found (S3N-P Fig).

To confirm that Shh activity was decreased by GPR17-mediated signalling, we performed a reporter assay to measure Gli activity. We prepared pools of explants transfected with the *GBS-Luc* reporter construct, which harboured the luciferase gene driven by the Gli binding sequence (GBS). The luciferase activity was measured after 24 hours. As the result, Gli activity was significantly upregulated by Shh^H^ (Fig 3L lanes 1 and 2). However, when MDL29951 was added along with Shh^H^, the activity was reduced to a level close to that of explants cultured with the Shh^L^ (more than four pools of explants in each condition; Fig 3L lanes 2-4). These data confirmed that the Gli activity was perturbed by GPR17 and its related signals.

We attempted to further investigate if the activation of the GPR17-related signal correlated with the elevation of the intracellular cAMP level, by assaying intermediate neural explants. Consistent with the previous observation, the cAMP level in the explants treated with Shh for 24 hours showed a lower cAMP level than in the control explants (Fig 3M, lanes 1 and 2) [18,21,39]. However, in the explants treated with MDL29951 in combination with Shh, the cAMP level was restored (Fig 3M, lanes 1 and 3). The observations suggested that MDL29951 perturbed the Shh-mediated decrease in the cAMP level.

Together, these findings suggested that the GPR17-mediated signalling pathway negatively regulates Shh activity in the context of neural tube pattern formation through the upregulation of the intracellular cAMP level.

### Perturbation of GPR17 expression causes aberrant expansion of ventral progenitor domains

To further investigate the functions of GPR17 in the development of neural tube pattern formation, we employed a loss-of-function approach. For this purpose, we designed a *siRNA* and a *shRNA* targeting GPR17 (*si-GPR17* and *sh-GPR17*), and carried out *in ovo* electroporation into the neural tube.

We first electroporated *si-GPR17* into neural tubes at HH stage 11. At 48 hpt, we observed aberrant expansion of the ventral neural domains characterised by Olig2, Nkx2.2 and FoxA2 (8 embryos for *si-control*; 10 embryos for *si-GPR17*; Fig 4A-4F”). Moreover, probably because the low level expression of *GPR17* was perturbed by *si-GPR17*, the area positive for Pax7 expressed in the dorsal progenitor domains was diminished (Fig 4G-4H”). This finding suggested that GPR17 *per se* is a negative regulator of the Shh signal, and that the expansion of ventral neural regions was caused by perturbation of *GPR17* expression. This aberrant expression was abolished by co-electroporation of mouse *GPR17*, which is not targeted by *si-GPR17* (6 embryos for each; S4D-S4F” Fig), suggesting that the phenotype observed in *si-GPR17* transfection was due to downregulation of *GPR17* expression. Furthermore, co-electroporation of *GPR161*, another negative regulator of the Shh signalling pathway [24], did not abolish the phenotypes caused by *si-GPR17*, confirming the specificity of *si-GPR17* (more than 6 embryos for each; S4G-S4I” Fig). The expansion of ventral identity was also observed when *sh-GPR17*, another knockdown DNA-based construct, was electroporated into neural tubes (more than eight embryos for each; S5 Fig), confirming the observations made with *si-GPR17*. Thus, GPR17 was demonstrated to be a negative regulator of the intracellular Shh signalling, and is essential for proper pattern formation in the neural tube.

**Figure 4:**
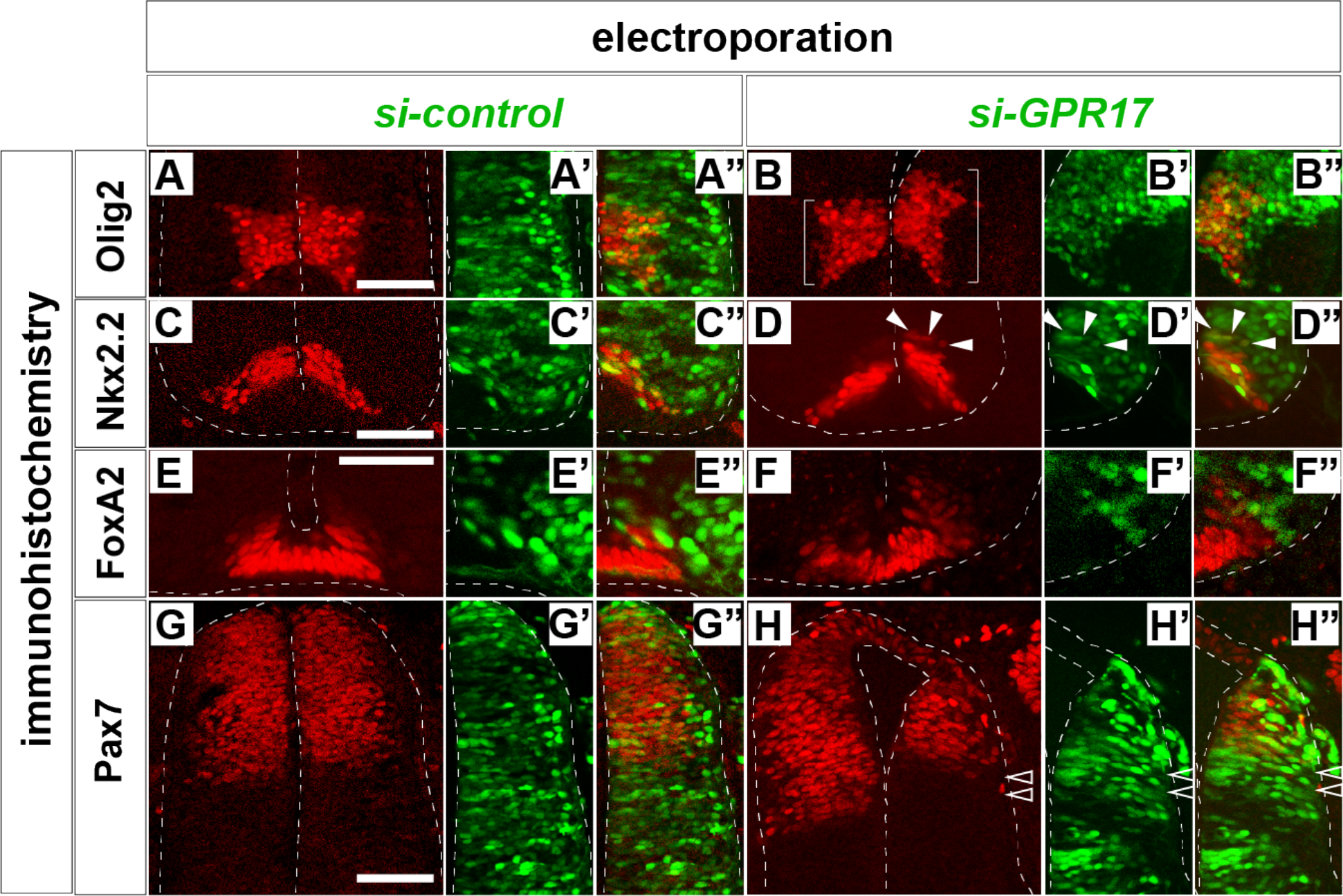
Knockdown of GPR17 affects dorsal-ventral pattern formation of the neural tube. The neural tube was electroporated with *si-control* (**A**-**A”**,**C**-**C”**,**E**-**E”**,**G**-**G”**) or *si-GPR17* (**B**-**B”**,**DD”**,**F**-**F”**,**H**-**H”**) at HH stage 12 on the right side of the neural tube, and incubated for 48 hours until the embryos reached HH stage 24. Sections of the neural tube were analysed with antibodies against Olig2 (**A**-**B”**), Nkx2.2 (**C**-**D”**), FoxA2 (**E**-**F”**) or Pax7 (**G**-**H”**). Scale Bar in **A** for **A-B”**, **C** for **C**-**D”**, **E** for **E**-**F”**, **G** for **G**-**H”** = 50 μm. Expanded and reduced areas are indicated by brackets, white arrowheads and outlined arrowheads, respectively.

### GPR17 is essential for dynamic control of Shh activity

Given that expression of *GPR17* is induced by Shh, and GPR17 negatively regulates the Shh signal activity at the intracellular level (Fig 1B, 1D, 2E, 3E-3L), we reasoned that GPR17 is involved in temporal regulation of Gli activity [13-15]. To test this hypothesis, we analysed the role of GPR17 in temporal regulation of the Gli activity.

First, to confirm that the intracellular Shh signal activity was aberrantly upregulated, we prepared intermediate neural explants electroporated with *si-GPR17*, and then treated them with Shh^L^. Although Olig2-expressing cells were predominant in the *si-control* electroporated explants (5 areas; Fig 5A, 5B, 5E, 5F, 5I), a significantly higher population of Nkx2.2-positive cells appeared at 24 hours in the *si-GPR17* electroporated explants (5 areas; Fig 5C, 5D, 5G, 5H, 5I), suggesting that the intracellular Shh signal activity was upregulated when *GPR17* expression was reduced. This observation was supported by the effect of treatment with pranlukast, a chemical antagonist of GPR17 [35,40]; explants treated with 10 μM pranlukast in combination with Shh^L^ tended to contain larger proportions of Nkx2.2-positive cells than controls, suggesting that perturbation of GPR17 caused the more ventral identity (10 areas each; S6A-S6G Fig).

**Figure 5:**
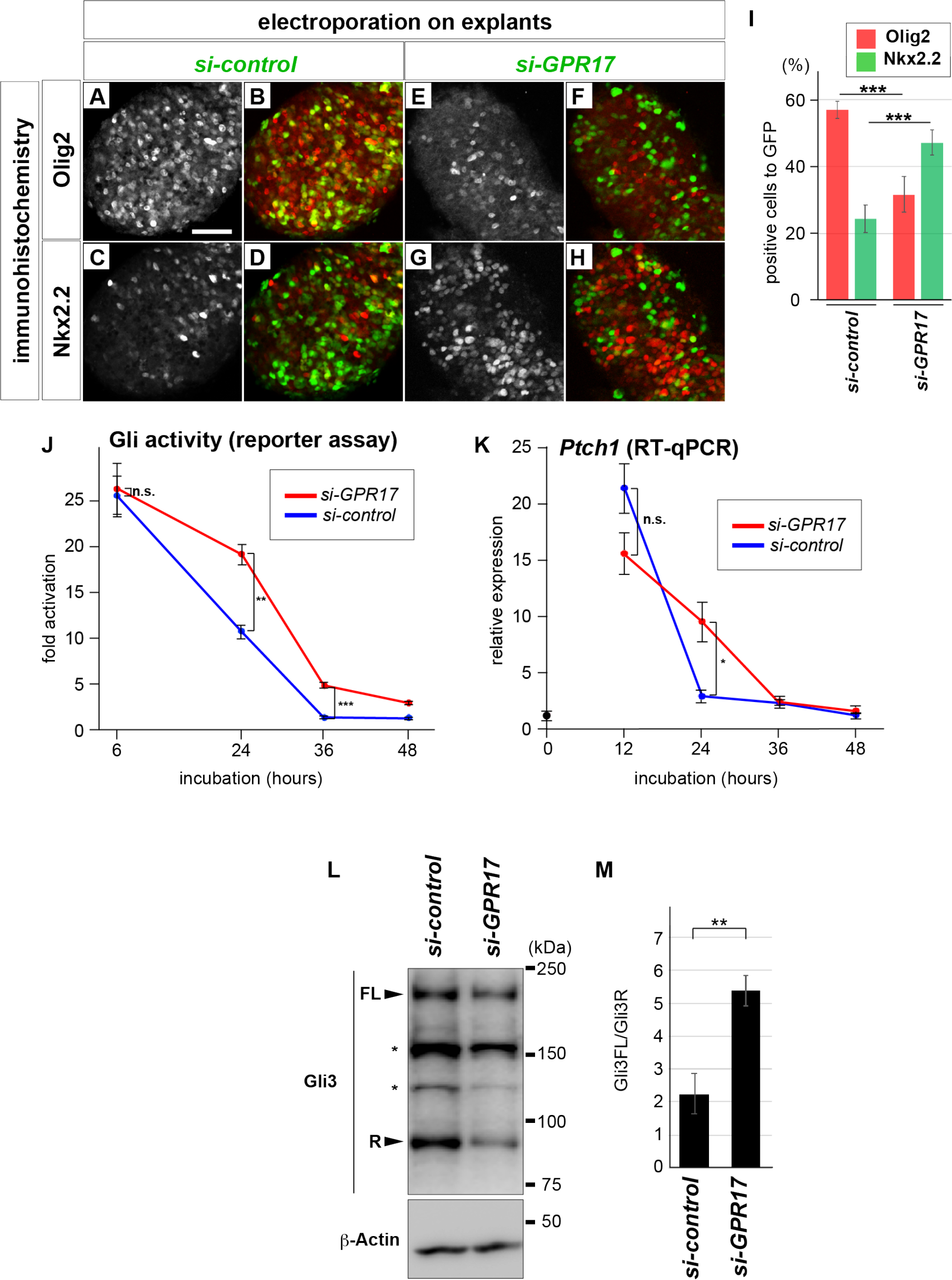
GPR17 controls temporal Gli activity. (**A**-**H**) Knockdown of *GPR17* induces more ventral identity. Intermediate neural explants electroporated with *si-control* (**A**-**D**) or *si-GPR17* (**E**-**H**) were treated with Shh^L^ for 24 hours, and expression was analysed with antibodies against Olig2 (**A**,**B**,**E**,**F**) or Nkx2.2 (**C**,**D**,**G**,**H**) and GFP (**B**,**D**,**F**,**H**). Merged images are shown in (**B**,**D**,**F**,**H**). (**I**) Quantitative data for (**A**-**H**). (**J**) feedback of Gli activity is partially perturbed by knockdown of GPR17. Explants electroporated with *si-control* (blue line) or *si-GPR17* (red line) together with GBS-Luc were incubated with Shh^L^ for the indicated time points, and luciferase activity was measured. (**K**) Prolonged expression of *Ptch1* in the intermediate neural explants treated with Shh^L^. *si-control-* (blue line) or *si-GPR17-* (red line) electroporated explants were treated with Shh^L^ and the *Ptch1* mRNA level was analysed in the samples harvested every 12 hours. (**L**,**M**) Suppression of GPR17 expression changes the ratio of the Gli3FL and Gli3R. Embryos were electroporated with either si-control or si-GPR17 on both sides of the neural tube, and then isolated after 24 hours. Neural tubes were manually isolated and analysed by western blotting with the Gli3 antibody (**L**). (**M**) Quantitative data from four independent experiments. Scale Bar in **A** for **A**-**H** = 50 μm. Expanded and reduced areas are indicated by white arrowheads and outlined arrowheads, respectively.

We next sought to analyse dynamic control of the intracellular Shh signal activity during ventral neural differentiation. For this purpose, we performed luciferase reporter assays using *GBS-Luc* at various time points. We prepared explants electroporated with *GBS-Luc* with either *si-control* or *si-GPR17*, and then measured Gli activity at a series of time points from 6 to 48 hours after the initiation of the culture. In *si-control*-electroporated explants treated with Shh^L^, Shh activity gradually decreased over time, peaking at 6 hours and becoming undetectable by the 48 hour time point (more than 3 explants per point; Fig 5J, blue line), consistent with previous reports [13,14]. On the other hand, in explants with *si-GPR17*, Gli activity at 6 hours was comparable to that of the *si-control* explants, but the activity was significantly higher at 24 hours and still detectable at 36 hours (Fig 5J, red line).

We next attempted to confirm that the dynamic Gli activity reflects the gene expression. As the *Ptch1* and *Gli1* genes are direct targets of the Shh signal, their expression levels correspond to the Gli activity [17]. We therefore cultured the explants electroporated with *si-control* or *si-GPR17* in the presence of Shh^L^ for different time periods, and measured the expression levels of *Ptch1* and *Gli1* by RT-qPCR. As the result, the expression peaked at 12 hours and then decreased quickly at 24 hours in *si-control* electroporated explants (Fig 5K and S6H Fig, blue lines). However, in the *si-GPR17-*electroporated explants, while the expression peak at 12 hours was comparable, the decrease was delayed, and the expression level was significantly higher at 24 hours (Fig 5K and S6H Fig, red lines). This result supports the idea that the dynamic Gli activity was affected, at least partially, by the perturbation of the GPR17 expression.

In addition, we calculated the ratio of Gli3FL and Gli3R in the neural tube. For this purpose, we prepared isolated neural tubes that had been electroporated with either *si-control* or *si-GPR17*, and analysed the forms of Gli3 by western blotting. We found that the ratio of Gli3FL over Gli3R was significantly higher in *si-GPR17*-electroporated neural tube than in the control (4 experiments; Fig 5L and 5M) [18].

Together, these findings indicate that GPR17 is an essential upstream factor that controls the dynamic change between the full-length and repressor forms of Gli3, and thus regulates the temporal change in the intracellular Shh signalling activity.

### GPR17 affects cell fate determination in neural differentiation from mouse ES cells

We finally assessed whether GPR17 functions are conserved in different organisms by investigating its expression and functions in aspects of mouse development.

We first analysed the expression pattern of GPR17 in the mouse spinal cord. At embryonic day 10.5 (e10.5) when neural tube pattern formation is taking place, expression was evident in the ventral region (S7A Fig) with the highest expression in the Olig2-positive cells (S7A, S7a’, S7a” Fig), with marginal expression in the dorsal area (S7A Fig). At e11.5, the expression was more ventrally restricted to the Nkx2.2-positive region (S7B, S7b’, S7b” Fig), with the neuronal expression (S7B Fig). This expression pattern was almost the same as in the developing spinal cord of chicks, regardless of some species-specific expression.

Next we investigated the requirement of GPR17 in the neural fate determination. For this analysis, we examined the directed neural differentiation of the embryonic stem (ES) cells, because the effect of treatments on gene expression are easily evaluated in this system. Mouse ES cells were differentiated into neural subtypes of pMN, p3 and floor plate using different combinations of retinoic acid (RA) and SAG, as described previously (S8A Fig) [41].

First, we evaluated GPR17 expression by RT-qPCR in each neural subtype. The results revealed significantly higher GPR17 expression in pMN and p3 cells than in FP 5 days after the start of differentiation (day 5) (3 experiments; Fig 6A), suggesting that expression can be recapitulated in neural cells differentiated from ES cells.

**Figure 6:**
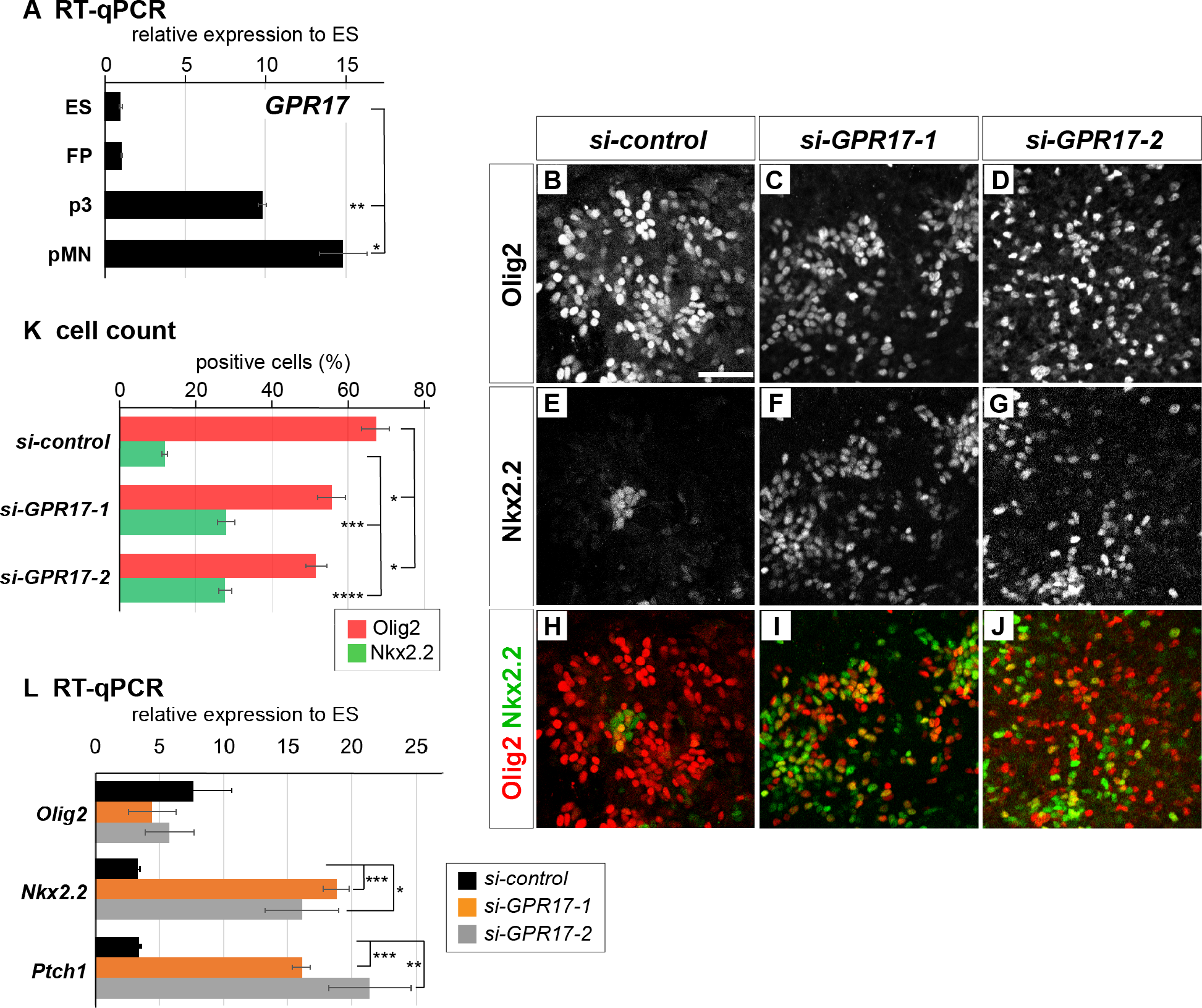
GPR17 is required for proper directed neural differentiation from mouse embryonic stem cells. (**A**) GPR17 expression level is subtype-specific. ES cells were differentiated according to the protocol described in **S8A Fig.**, and the expression levels of GPR17 in each sample were analysed by RTqPCR. (**B**-**J**) Directed neural differentiation from ES cells. ES cells transfected with *si-control* (**B**,**E**,**H**), *si-GPR17-1* (**C**,**F**,**I**) or *si-GPR17-2* (**D**,**G**,**J**) were differentiated according to the protocol (i) (**S8A Fig**), which mainly yields pMN cells, and analysed with antibodies against Olig2 (**B**-**D**) and Nkx2.2 (**E**-**G**). Merged images are shown in (**H**-**J**). (**K**) Quantification of Olig2 and Nkx2.2 expressing cells over DAPI positive cells. (**L**) Expression levels of *Olig2*, *Nkx2.2* and *Ptch1* were analysed by RT-qPCR. Scale bar in **B** for **B**-**J** = 50 μm.

Next, we sought to analyse the function of GPR17 in neural differentiation and subtype determination, and transfected two different *si-RNAs* (*si-GPR17-1* and *si-GPR17-2*) targeting *GPR17* in ES cells. The efficient knockdown of *GPR17* expression was validated by RT-qPCR (3 experiments; S8B Fig).

Under these conditions, ES cells were differentiated into pMN cells (S8A Fig), and gene expression was evaluated on day 5. In the pMN condition, the majority of cells expressed Olig2, with a small subset of Nkx2.2-positive cells, as demonstrated by immunohistochemistry (3 experiments; Fig 6B, 6E, 6H, 6K). By contrast, treatment with *si-GPR17* increased the Nkx2.2-expressing cells (Fig 6E-6K). This observation suggests that *si-GPR17*-treated cells became more susceptible to the Shh ligand and tended to differentiate into the more ventral identity of p3 (S8C-S8E Fig), for which a higher Gli activity was required [41].

We next verified this tendency by RT-qPCR. In cells treated with *si-GPR17s*, the Nkx2.2 expression was higher than in *si-control*-transfected cells at day 5, confirming the results obtained by the immunohistochemistry (3 experiments; Fig 6L). Furthermore, the expression level of *Ptch1*, a target gene of the Shh signal, was higher in *si-GPR17* cells than in the controls (3 experiments; Fig 6L). These results were consistent with the findings obtained in the analyses of chick embryos.

These findings suggest that mouse GPR17 is expressed in the pMN and p3 cells differentiated from ES cells as in the chick embryos, and functions as a modulator for the ventral identities of the neural cells.

## Discussion

### GPR17 is a negative regulator of the Shh signalling pathway and affects ventral pattern formation of the neural tube

In this study, we isolated one of the GPCRs GPR17, and characterised its role in neural development. Although its expression is induced by Shh, GPR17 negatively regulates the Shh signal, thereby affecting the dynamicity of the Gli activity [13,14]. Negative regulation of the intracellular Shh signal by GPR17 is conserved in the mouse, as demonstrated by a system involving neural differentiation of ES cells (Fig 6).

GPR17 was initially recognised as one of the genes that respond to neural tube injury, brain damage or pathological situations [42-44]. To reveal the essential roles of GPR17 in the body, *GPR17-*knockout mice were generated. The analysis consequently elucidated that GPR17 is a molecular timer for oligodendrocyte differentiation [44]; the deficiency of *GPR17*-gene caused the earlier differentiation and excessive production of oligodendrocyte cells [27]. Subsequently, the function in the oligodendrocyte development and the demyelination and remyelination have been emphasised. GPR17 expression is gradually upregulated during the demyelination induced by the glial toxin Lysolecithin [45]. Conversely, the antagonist supporting GPR17 increases the oligodendrocyte cell number [46]. Moreover, the proliferation of the Olig2-expressing cells is encouraged in the situation where GPR17 is attenuated [45]. Given that oligodendrocyte differentiation is supported by Shh [47], the phenotypes caused by the blockade of GPR17 expression or functions could be due to hyper-activation of Gli activity. Therefore, the notion we propose here - GPR17 is a negative regulator for Shh signalling pathway, especially for the Gli activity - is consistent with the previous findings obtained using this *GPR17*-deficient mouse [27].

Although GPR17 has been suggested to be a receptor for the uracil nucleotides and Cysteinyl leukotrienes (cysLTs) [42,48], the actual ligand(s) for GPR17 working in the developmental contexts have not been identified. In this study, while we did not find an explicit effect of the overexpression of GPR17 *in vivo* (S4A-S4C” Fig), it is still possible that GPR17 may be activated by the endogenous ligands and that the identification of such ligands is one of the important questions to be addressed in the future.

In this study, we utilised MDL29951 to experimentally activate GPR17 (Fig 3F, 3H, 3J, 3K-3M, S3E-S3E”, S3J-S3J” Fig). While MDL29951 has been recognised as an agonist of GPR17 [35] and was actually used to analyse the function of GPR17 [28], MDL29951 has also been recognised as an antagonist of the N-methyl-D-aspartate (NMDA) receptor [49,50]. However, as we demonstrated, MDL29951 synergistically acted with GPR17 (S3 Fig) and it is highly likely that MDL29951 works in concert with GPR17 to determine the pattern formation and fate determination of the developing neural cells.

In the chick embryos, our *in situ* hybridisation analysis detected the expression of *GPR17* at the earlier stages than the onset of gliogenesis, which contributed to neural tube pattern formation (Fig 1A, 1B, 1D). Also in the mouse embryos, the immunohistochemistry analysis revealed the *GPR17* expression at e10.5 at the ventral region of the spinal cord (S7A Fig). By contrast, in a previous report [27], GPR17 expression is firstly recognised at e14.5 in the mouse brain using reporter gene expression. The difference between our data and the previous findings is presumably caused by the protein stability of GPR17 or the difference in the detection methods.

While our analysis revealed the essential roles of GPR17 on the dorsal-ventral pattern formation of the spinal cord in chick, the effect in the mouse neural tube has been unknown, and at least, does not seem to be critical [27,45,46], as *GPR17*-deficient mice are viable. This is probably due to the redundant roles of the multiple GPCRs expressed in the neural tube. The expression of *Adenylyl cyclase 5* (*AC5*), which catalases the dissociation of ATP to make cAMP [20,21], was found to be induced by Shh both in chick and mouse neural explants (S9 Fig). Thus, it is possible that GPR17 and AC5 have redundant roles in the neural tube development, and that the dependency on multiple GPCRs may differ among different species; the chick neural tube mainly depends on *GPR17* in the pattern formation while the mice depend also on the other genes.

GPCRs can bind to different types of Gα proteins, including Gα_i_, Gα_s_, Gα_q_ and Gα_12,13_ [23]. Among these Gα proteins, Gα_q_ and Gα_s_ can increase the intracellular cAMP level, whereas Gα_i_ proteins mostly exert an inhibitory effect, and GPR17 can bind all types of G-proteins [35]. During the remyelination, GPR17 binds to Gα_i_ and decreases the cAMP level [28,36,51,52]. By contrast, in our experiments GPR17 rather upregulated the cAMP level, presumably by binding to Gα_q_ and/or Gα_s_, suggesting that it has diverse and cell type-specific functions.

Concerning the relationships between GPCR and Shh signal, two GPCRs, GPR161 and GPR175, have been well characterised [22,24,25]. GPR161 is an orphan receptor that increases the intracellular cAMP level and has a negative effect on the ventral neural pattern formation. Within the cell, GPR161 is localised at the ciliary shaft and is involved in local calcium uptake when Shh protein arrives at the cilia [21,24]. GPR175, which localises to non-ciliary cell membrane, is translocated to the cilia when cells are exposed to Shh [25]. By contrast, GPR17 is localised throughout in the cells (Fig 2B), and its localisation is not affected by Shh signals (Fig 2B-2D’), suggesting that the roles of GPR17 are distinct from those of these other GPCRs. At the tissue level, expression of GPR161 is ubiquitous and that of GPR175 is unknown, whereas GPR17 expression is explicitly upregulated by Shh. The relationship among multiple GPCRs in regulation of the intracellular Shh signal is an intriguing issue to be addressed in the future.

### GPR17 constitutes a negative feedback loop of the Shh signalling system

A historical model suggested that a morphogen can produce a number of cell types depending on the thresholds of the signal concentration [53]. This hypothesis relies on the supposition that signal concentration corresponds completely to the intracellular signal intensity. However, recent studies suggest that the temporal change in signal activity, so-called temporal adaptation, is important for cell fate determination and correct pattern formation [1,13,14]. The results of this study reveal a novel negative feedback loop constructed by a GPCR through regulation of the intracellular cAMP level and processing of Gli transcription factors. Recent mathematical modelling suggests the existence of negative regulators induced by Shh [17], and our findings regarding GPR17 are in good agreement with this hypothesis.

Although Shh induces the *GPR17* expression (Fig 1D), *GPR17* does not seem to be a direct target gene of Gli transcription factors, as *GBS* has not been identified in the flanking regions of the *GPR17* locus [41]. Instead, Olig2 is the direct regulator of *GPR17* expression [28]. Consistent with this finding, overexpression of Olig2 in neural explants induces *GPR17* expression (Fig 1E). This could explain why the negative feedback regulation is cell type-specific and does not take place in the NIH3T3 cells treated with the Shh ligand [17]; Olig2 expression is induced only in the neural progenitor cells, but not in NIH3T3 cells (AY, AH, NS; unpublished observation).

While it was shown that Olig2 directly induces the *GPR17* expression [28], Olig2, together with its related factor Olig1 [54], has been shown to be a transcriptional repressor [55], and the activator type of Olig2 (Olig2^DBD^-VP16) [54] works as an antimorphic mutant to repress the *GPR17* expression (S10 Fig). This apparent discrepancy could be explained by the existence of cofactor(s) or other transcription factor(s), whose expression depends on Olig2. Moreover, it is possible that the other factors are involved in the *GPR17* expression [56], as the *GPR17* expression is evident in the domains where Olig2 is not expressed (Fig 1B), and the overexpression of Olig2^DBD^-VP16 only partially block the *GPR17* expression (S10B Fig).

In a developmental context, negative feedback regulation confers diversity of cell types and robustness of pattern formation [14]. To date, multiple negative feedback systems for the Gli activity have been identified [17]. For instance, the *Ptch1* gene, whose product negatively regulates the intracellular Shh signal activity, is a direct target of Shh (Fig 7) [13]. Moreover, expression of the transcriptional mediator Gli2 decreases over time during neural development, although the underlying regulatory mechanisms remain elusive [17]. In combination with the existence of GPR17 as this study demonstrated (Fig 7), multiple negative feedback systems could cause differences in the half-life of the signal in each progenitor cell, allowing diverse cell types to be produced by a single morphogen, Shh. Moreover, during formation of the morphogen gradient, uncertainties and fluctuations may arise. This noise in the signal is constantly modulated by negative feedback, and consequently the reliability and reproducibility of pattern formation can be incarnated [57,58]. Therefore, the negative feedback loop formed by Shh and GPR17 is important not only for overall neural tube pattern formation, but also for fine tuning of signal intensity, and consequently establishes the quantitative balance among different cell types during neural and neuronal differentiation. Further studies, including analysis of the temporal change of intracellular cAMP level, will reveal the significance of GPR17 in negative feedback of the Shh signal.

**Figure 7:**
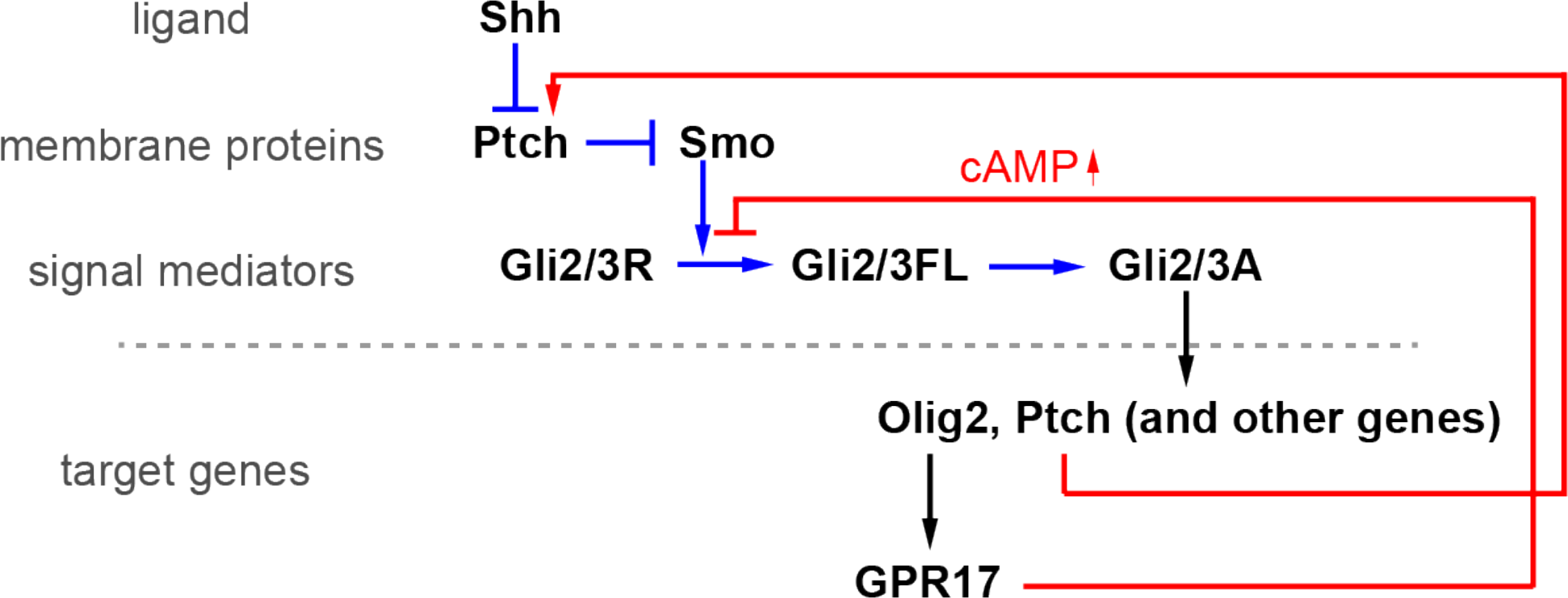
A working model for the negative regulation of the intracellular Shh signaling pathway. The binding of Shh (ligand) to Ptch (a membrane protein) abrogates its repressive effect on Smo, and Gli2/3 are thereby converted from their repressor forms (Gli2/3R) to the full-length forms (Gli2/3FL). Gli2/3FL are further modified to their active forms (Gli2/3A) and are translocated to the nucleus where the target genes, including *Olig2* and *Ptch*, are induced. It has been shown that the accumulation of *Ptch* transcript results in the perturbation of the Shh signal transduction. In addition, as demonstrated in our study, Olig2 induces the *GPR17* gene expression, and the conversion of Gli2/3R to Gli2/3FL is inhibited by the upregulation of cAMP. The positive and negative regulations are indicated by blue and red lines, respectively. The transcriptional regulation is indicated by black lines.

## Materials and Methods

### Isolation of GPR17

GPCRs that can couple with Gα_q_ or Gα_s_ were identified by referring to the website (http://gpcrdb.org), and qPCR primers were designed against the corresponding genes (S1 Table). Relative expression levels were analysed by RT-qPCR in neural explants treated or untreated with Shh^H^ (S1 Fig). NCBI Gene IDs for chicken and mouse *GPR17* are 769024 and 574402, respectively.

### Embryos, electroporation and expression analysis

All animal experiments were performed with the approval of the Animal Welfare and Ethical Review Panel of Nara Institute of Science and Technology (approval numbers 1636 and 1810 for chicken and mouse experiments, respectively). Chicken eggs were purchased from the Shiroyama Keien farm (Kanagawa prefecture, Japan) and the Yamagishi farm (Wakayama prefecture, Japan), and the developmental stages were evaluated by the Hamburger and Hamilton’s criteria [59]. *In ovo* electroporation of chick embryos was carried out with an ECM830 electroporator (BTX), and embryos were incubated at 38°C for the indicated times. Overexpression in the chick embryos was performed with the constructs subcloned into the *pCIG* expression plasmid, which contains an IRES (internal ribosomal entry site) -GFP gene downstream of the gene of interest [60]. *pCIG-Olig2^DBD^-VP16* was constructed by fusing the DNA-binding region of Olig2 with the transactivation domain of the viral protein VP16, as described previously [55]. Embryos were fixed with 4% paraformaldehyde, subsequently incubated with 15% sucrose for cryoprotection and embedded in 7.5% gelatine (Sigma). Cryosections were cut at 14 μm increments and analysed with immunohistochemistry or in situ hybridisation. The sections from at least five independent embryos were analysed, and the number of the embryos analysed were indicated in the text. For loss-of-function experiments, *si-GPR17* (GGAACAGAGUGGAGAAACACCUG(dA)(dA) (sense) and UUCAGGUGUUUCUCCACUCUGUUCCCA (antisense)) or *si-Luciferase* (ACUGAGACUACAUCAGCUAUUCU(dG)(dA) (sense) and UCAGAAUAGCUGAUGUAGUCUCAGUGA (antisense), as a negative control; Eurofin) was electroporated with the *pCIG* vector, which encodes *GFP* as a tracer. For gene silencing with short-hairpin constructs, AAGAGACACACCTCGAGAATG (chicken *GPR17*) or AATGTATTGGCCTGTATTAGC (control) was subcloned into the *pRFP-RNAiC* vector [61].

DIG-labelled (Roche) complementary RNA probe was synthesised using RNA polymerase (Promega). *In situ* hybridisation on slides was performed as described previously [62]. For western blotting, neural tube tissues electroporated with *siRNAs* were manually isolated from embryos and dissociated in TN Buffer (150 mM NaCl, 5 mM KCl, 0.5% NP-40 detergent, 10 mM Tris-HCl, pH 7.8) with protease inhibitor cocktail (Roche).

Immunohistochemistry was performed as described previously [62]. Primary antibodies against the following proteins were used in this study: Olig2 (rabbit, Millipore AB9610), Nkx2.2 (mouse, DSHB 74.5A5; rabbit, Abcam ab19077), Islet1/2 (DSHB 39.4D5), Pax7 (DSHB), FoxA2 (goat, Santa Cruz sc-6554X), Sox2 (rabbit, Millipore AB5603), GPR17 (goat, Abcam ab106781 for mouse embryos; rabbit, Sigma SAB4501250 for cells), GFP (sheep, AbD Serotec 4745-1051), Gli3 (goat, R&D AF3690), β-Actin (rabbit, Abcam ab8227), and γ-acetylated tubulin (mouse, Sigma T7451). In the western blots with this Gli3 antibody, we always found two unexpected bands located at approximately 160 and 110 kDa, whose intensities accompanied that of Gli3, even though these bands were not Gli3. For GPR17 immunochemistry in mouse embryonic sections, antigenic retrieval was required; slides were boiled for 1 minute in 10 mM of citric acid (pH 6.0) solution prior to the incubation with the primary antibody. Fluorophore- and HRP-conjugated secondary antibodies were purchased from Jackson Laboratory and Cell Signaling Technology, respectively.

### New culture and intermediate neural explants

New culture was performed as described previously [37]. Culture plates containing albumen and agarose were prepared with the final concentration of 100 μM of MDL29951 or with dimethylsulfoxide (DMSO) (control). Chick embryos were taken from the eggs at HH stage 12 with a ring of filter paper, and put on the culture plates with the ventral side up. Embryos were cultured in the 38°C incubator for 36 hours, and the plates were kept humid with Hanks’ Balanced Salt solution (1xHBSS; Sigma-Aldrich).

Chick intermediate neural explants were prepared as described previously [62]. Briefly, HH stage 9 embryos were isolated from the eggs and maintained in L-15 medium (Thermo Fisher Scientific). After treated with dispase II (Sigma), the intermediate region of the neural plate at the preneural tube level [26] were manually excised from the embryos and cultured in DMEM/F-12 medium (Thermo Fisher Scientific) supplemented with Mito+Serum Extender (Sigma) and Penicillin/Streptomycin/Glutamine (Wako). Quantitation of positive cells was performed on at least five areas, each of which contained approximately 200-250 cells, randomly chosen from independent explants. For RT-qPCR analyses, RNA was extracted from the pools of 15 chick intermediate neural explants by using PicoPure RNA extraction kit (Thermo Fisher Scientific). Each extraction gave a range of 500 ng (12-hour culture) to 1.5 μg (48-hour culture) of total RNA, as measured by Nanodrop (Thermo Fisher Scientific). Complementary DNAs (cDNAs) were synthesised by PrimeTaq reverse transcriptase (TaKaRa), and qPCR was performed by using LightCycler 96 (Roche). At least two independent pools of explants were analysed in each experiment, and each gene expression level was normalised with the *GAPDH* expression.

Mouse neural explants (S9 Fig) were prepared from the e8.5 embryos and were cultured in the same way as in the chick explants [15], except that the culture medium contained N2 and B27 supplements (Thermo Fischer Scientific).

### Cell culture, transfection and selection

NIH3T3 fibroblast cells were maintained in Dulbecco’s modified Eagle’s medium /F12 medium (Wako) containing 10% New-Born Calf Serum (MP Biomedicals). Lipofectamine 3000 (Invitrogen) was used for transfection. To obtain explicit evidence of the effects on the overexpression of the genes in **Fig 2E**-**2I**, the transfected cells were selected for 48 hours with 500 μg/ml of G418 (Invitrogen) from one day after the *pcDNA3*-based expression plasmids were transfected, and were used for assays.

### Monitoring the Gli activity and the intracellular cAMP levels

The *GBS-Nano-Luc* reporter construct containing the *GBS* was constructed by subcloning of the octameric *GBS* [63] to the pNL3.2 vector (Promega), and used to monitor Gli activity. *pCS2-firefly luciferase* was used for an internal control. For the luciferase assays on chick neural explants in Fig 3L and Fig 5J, *GBS-Nano-Luc* and *pCS2-fiefly-luciferase* were electroporated at the preneural tube region of the caudal part of the chick embryos at HH stage 9- [26]. *si-RNAs* were co-electroporated if necessary. The explants were then prepared and were cultured with the indicated conditions. The relative luciferase activities were compared to those cultured without Shh at every time point. 7-8 explants were grouped for each measurement, and four to 10 groups of explants were measured in one condition.

*CREB-Luc* (luciferase gene driven by the cAMP-responsive element-binding region) was a kind gift from Prof. Itoh [64] and *pRL-CMV* (Promega) was used for an internal control. The measurement of the chemiluminescence was performed by using the plate reader Tristar2 (Berthold Technologies).

For the cAMP assay in the NIH3T3 cells (Fig 2H), the cells were transfected with the indicated plasmids and the transfected cells were selected for 24 hours. 3-isobutyl-1-methylxanthine (IBMX) was added at 1 μM in the last 30 minutes before cells were harvested. For the cAMP assay in the neural explants (Fig 3I), 15 explants were prepared for each condition and cultured for 24 hours. IBMX was added in the last 1 hour. The cAMP assay was performed using DetectX high sensitivity direct cAMP Chemiluminescent Immunoassay Kit (Arbor Assays). The protein concentrations of the cell lysates were measured by CBB protein assay solution (Nacalai, Japan) and the measurement values were normalised.

### TUNEL assay

TUNEL assay was performed to detect the apoptotic cells. The neural tube sections or the explants were incubated with TdT transferase (Roche) and DIG-labelled dUTP (Merck Millipore), and the signals were visualised by anti-Digoxigenin-Rhodamine antibody (Sigma). Staurosporine [65] was used as a positive control at 1nM.

### Neural differentiation of mouse ES cells

The Sox1-GFP ES cell line was kindly provided by Prof. Smith, and maintenance and neural differentiation as a monolayer culture were performed as described previously [41,66,67]. Briefly, on the fibronectin/collagen-coated plates, approximately 1.5 × 10^4^ ES cells were seeded and cultured as monolayers in the differentiation medium [41] for the initial 3 days. For pMN differentiation, cells were added with 300 nM RA (Sigma) on day 3.0 and 50 nM SAG was added on day 3.5; and the cells were incubated for totally 5 days. For p3 differentiation, 30 nM RA and 500 nM SAG (Sigma) were added on day 3.0 and day 3.5, respectively. For FP differentiation, 500 nM SAG was added on day 3.0. Before differentiation was initiated, Stealth *siRNAs* (Invitrogen) (*si-GPR17-1*, UCGCCUGCUUCUACCUUCUGGACUU; *si-GPR17-2*, ACCGUUCAGUCUAUGUGCUUCACUA) or *si-control* (Invitrogen) were transfected twice at 24 hour intervals using Lipofectamine RNAiMAX (Invitrogen). For RT-qPCR analysis, RNA was extracted by RNeasy RNA extraction kit (QIAGEN). cDNA synthesis and qPCR were performed as in the chick neural explants.

### Images and data analysis

Images were collected by using AxioVision2 (for *in situ* hybridisation images; Zeiss), LSM 710 confocal microscope (for immunohistochemistry data; Zeiss), LAS4000 (for western blots; GE Healthcare), and signal intensities of the western blots were calculated by ImageJ. Images were processed by the software Photoshop (Adobe) and figures were arranged on Illustrator (Adobe). Statistical analysis was performed with Prism (GraphPad) by two-tailed t-test. Statistical data are presented as mean values ± s.e.m., and significance (*; p<0.05; **; p<0.01, ***; p<0.001, ****; p<0.0001) were indicated in each graph.

## Acknowledgements

The authors are grateful to Profs. Austin Smith and Hiroshi Itoh for materials, Dr. James Briscoe for comments on the manuscript, Dr. Yuichi Sakumura for advice, Prof. Takashi Toda for encouragement, and all laboratory members for support and valuable discussion.

## Data Availability

All relevant data are within the text, figures, and their Supporting Information files.

## Author Contributions

Conceived the project: NS. Performed the experiments, analysed the data: AY, AH, MK, NS. Providing essential materials and advice: MMT, TK. Writing the manuscript: NS.

## Supporting Information Legends

**S1 Fig. RT-qPCR screening for Gα_q_- and Gα_s_-coupled GPCRs responsive to Shh.**Neural explants were treated with Shh^H^ for 24 hours, and the expression of each gene was analysed by RT-qPCR. FoxA2 and Nkx2.2 were assayed as positive controls. Primer sequences used for the qPCR are listed in S1 Table.

**S2 Fig. Intermediate neural explants differentiate into the floor plate in Shh^H^, but into the motor neuron in Shh^L^, after 48 hours.**

(**A**) Schematic representation of progenitor domains and neuronal subtypes along the dorsal-ventral axis, along with the transcription factors expressed in the corresponding regions. (**B**) Islet-1 (green) and FoxA2 (red) are expressed in motor neurons and the floor plate region, respectively. (**C**-**H**) Shh^L^and Shh^H^ give rise to motor neurons and floor plate at 48 hours, respectively. Intermediate neural explants were treated with Shh^L^ (**C**,**E**,**G**) or Shh^H^ (**D**,**F**,**H**), and expression was analysed by immunohistochemistry using antibodies against Islet-1 (motor neuron: **B**,**C**,**D**,**G**,**H**; green) and FoxA2 (floor plate: **B**,**E**,**F**,**G**,**H**; red). Merged images are shown in (**G**,**H**). Scale Bar in **B**, in **C** for **C**-**H** = 50 μm

**S3 Fig. GPR17 in combination with its specific agonist MDL29951 affects the dorsal-ventral patterning of the neural tube.** Expression plasmids of control *GFP* (**A**-**A”**,**F**-**F”**), *SmoA1* + *control GFP* (**B**-**C”**,**G**-**H”**) or *SmoA1* + *GPR17* (**D**-**E”**,**I**-**J”**) were electroporated HH stage 12 and harvested at 48 hpt. During the incubation, vehicle (**A**-**B”**,**F**-**G”,D**-**D”**,**I**-**I”**) or 30 μM of MDL29951 (**C-C”**,**EE”**,**H-H”**,**J-J”)** was administered twice (at 12 hpt and 36 hpt) directly into the cavity of the neural tube. Immunohistochemistry was performed on the sections of neural tube with Nkx2.2 (**A**-**E”**) and Pax7 (**F**-**J”**) antibodies. The double-positive cells for Pax7 and GFP are indicated with white arrowheads (**J**-**J”**). The electroporated side of the neural tube is shown, and the ventral border of the endogenous Pax7-expressing area is indicated by yellow arrowheads (**F**-**J”**). Scale bar in **A** for **A**-**M”** = 50 μm. (**K**) Quantitative data for (**A**-**J’**). For Nkx2.2, the double-positive cells for GFP and ectopically positioned Nkx2.2 over the total GFP cells were indicated. For Pax7, the double-positive cells for GFP and Pax7 over the total GFP positive cells in the dorsal region were indicated. (**L**-**P**) MDL29951 does not induce apoptosis. TUNEL assay was performed on the neural tube sections electroporated with *SmoA1* (as in **B**-**B”** and **G**-**G”**; **L**-**L”**) or *SmoA1* + *GPR17* and subsequent treatment with MDL29951 (as in **E**-**E”** and **J**-**J”**; **M**-**M”**). (**N**-**P**) TUNEL analysis on explants. Chick neural explants treated with Shh^H^ (as in **Fig 3E, 3G, 3I**; **N**) or Shh^H^ + MDL29951 (as in **Fig 3F**, **3H**, **3J**; **O**) for 24 hours. Explants treated with 1 nM of staurosporine with Shh^H^ for 24 hours were used as a positive control (**P**). Scale bar in **N** for **N**-**P** = 50 μm.

**S4 Fig. GPR161 does not rescue the phenotype induced by *si-GPR17*, whereas mouse GPR17 does.**

*mGPR17* (**A**-**F**’) or *mGPR161* (**G**-**I**’) was electroporated in combination with *si-GPR17* (**D**-**I’**) at HH stage 12 on the right side of the neural tube, and embryos were incubated for 48 hours. Neural tube patterning was analysed with antibodies against Pax7 (**A**-**A**”,**D**-**D**”,**G**-**G**”), Olig2 (**B**-**B”**,**E**-**E”**,**H**-**H**”) and Nkx2.2 (**C**-**C**”,**F**-**F**”,**I**-**I**”). The ventral end of the Pax7 expression is indicated by arrowheads (**A**,**D**,**G**), and the reduced expression is indicated by outlined arrowhead (**G**). The expanded Olig2 and Nkx2.2 expression is indicated by brackets (**H**-**H**”) and arrowheads (**I**-**I**”), respectively. Scale Bar in **A** for **A**,**A**’,**B**,**B**’,**D**,**D**’,**E**,**E**’,**G**,**G**’,**H**,**H**’, in **C** for **C**,**C**’,**F**,**F**’,**J**,**J**’ = 50 μm

**S5 Fig.*sh-GPR17* induces a similar phenotype to *si-GPR17*.**

sh-control (**A**-**A”,C**-**C”,E**-**E”,G**-**G”**) or *sh-GPR17* (**B**-**B”,D**-**D”,F**-**F”,H**-**H”**) was electroporated on the right side of the neural tube at HH stage 12, and embryos were incubated for 48 hours. Neural tube patterning was analysed with antibodies against Pax7 (**A**-**B**”), Olig2 (**C**-**D**”), Nkx2.2 (**E**-**F**”) or FoxA2 (**G**-**H**”). The expanded and reduced expression is indicated by white arrowheads and outlined arrowheads, respectively. Scale bar in **A** for all images = 50 μm

**S6 Fig.*si-GPR17* induces a more ventral identity in neural differentiation.**

(**A**-**F**) Pranlukast, an antagonist of GPR17, elevated Shh activity. Neural explants were treated with ShhL for 24 hours along with DMSO (control; **A**,**C**,**E**) or 10 nM pranlukast (PLK), and then analysed using antibodies against Olig2 (**A**,**B**) or Nkx2.2 (**C**,**D**). Merged images are shown in (**E**,**F**). Quantitative data are shown in (**G**). Scale Bar in **A** for **A**-**F** = 50 μm. (**H**) *si-GPR17* causes the temporal expression of the *Gli1* gene. The analysis was performed as in **Fig 5K**, except the qPCR was performed with the *Gli1* primers.

**S7 Fig. GPR17 is expressed mainly in the ventral region in developing mouse spinal cord.**

Immunohistochemical analysis of GPR17 in the developing mouse spinal cord at embryonic day 10.5 (e10.5) (**A**) and e11.5(**B**). The positive areas for GPR17 are indicated by a bracket (**A**; ventral) and an arrowhead (**A**; dorsal) and arrowheads (**B**). The double-staining images with Olig2 (green: **a’**,**b’**) or with Nkx2.2 (green: **a”**,**b”**) are indicated. Scale bar = 50 μm (**A**,**a’**,**a”**,**b’**,**b”**) and 100 μ m (**B**).

**S8 Fig. Differentiation protocol of the ES cells and validation of *si-GPR17*.** (**A**) Protocol for differentiation of ES cells into pMN, p3 and FP cells. (**B**) The decrease in *GPR17* expression was validated by RT-qPCR. (**C**-**E**) Cells differentiated with the protocol (ii), which yields mainly p3 cells. Cells were analysed with antibodies against Olig2 (**C**) or Nkx2.2 (**D**). Merged image is shown in (**E**). Scale Bar in **C** for **C**-**E** = 50 μm.

**S9 Fig. *AC5* expression is induced by Shh.** Neural explants were prepared from chick (**A**) or mouse (**B**) embryos and were cultured for 24 hours without (black bar) or with Shh^H^ (red bar). The expression of *AC5* and *GPR17* was analysed by RT-qPCR.

**S10 Fig. The electroporation of dominant-negative Olig2 reduces the GPR17 expression.** *pCIG* (control vector conveying *GFP*; **A,A’**) or *pCIG-Olig2^DBD^-VP16* (**B,B’**) were electroporated at HH stage 12 and embryos were cultured for 48 hours. The expression of *GPR17* was analysed by *in situ* hybridisation (**A,B**), and the adjacent sections were analysed by immunohistochemistry with the GFP antibody (**A’,B’**). E.P.; electroporation, ISH; *in situ* hybridisation, IHC; immunohistochemistry.

**Supplementary Table 1 The PCR primers used in this study.**

